# Prediction of protein-protein interactions based on elastic net and deep forest

**DOI:** 10.1101/2020.04.23.058644

**Authors:** Bin Yu, Cheng Chen, Zhaomin Yu, Anjun Ma, Bingqiang Liu, Qin Ma

## Abstract

Prediction of protein-protein interactions (PPIs) helps to grasp molecular roots of disease. However, web-lab experiments to predict PPIs are limited and costly. Using machine-learning-based frameworks can not only automatically identify PPIs, but also provide new ideas for drug research and development from a promising alternative. We present a novel deep-forest-based method for PPIs prediction. First, pseudo amino acid composition (PAAC), autocorrelation descriptor (Auto), multivariate mutual information (MMI), composition-transition-distribution (CTD), and amino acid composition PSSM (AAC-PSSM), and dipeptide composition PSSM (DPC-PSSM) are adopted to extract and construct the pattern of PPIs. Secondly, elastic net is utilized to optimize the initial feature vectors and boost the predictive performance. Finally, GcForest-PPI model based on deep forest is built up. Benchmark experiments reveal that the accuracy values of *Saccharomyces cerevisiae* and *Helicobacter pylori* are 95.44% and 89.26%. We also apply GcForest-PPI on independent test sets and CD9-core network, crossover network, and cancer-specific network. The evaluation shows that GcForest-PPI can boost the prediction accuracy, complement experiments and improve drug discovery. The datasets and code of GcForest-PPI could be downloaded at https://github.com/QUST-AIBBDRC/GcForest-PPI/.

## 1. Introduction

The study of the protein-protein interactions (PPIs) of molecular mechanisms is essential (Alberts, 1998; Amar, Hait, Izraeli & Shamir, 2015; Schadt, 2009). The disorder of the PPI network structure can cause abnormalities in cell life activities. Because of the progress of high-throughput technologies, lots of PPIs via web-lab experimental verification have emerged. Multiple PPIs sources lead to the generation of PPIs databases, containing the DIP (Xenarios, et al., 2002), and HPRD (Peri, Navarro, Amanchy & Kristiansen, 2003). The detection of PPIs relied on computational methods could reduce the web-lab limitations and an effective, accurate, useful machine learning algorithm can predict large scale PPIs.

Several genomic features have been used in PPIs prediction based on machine learning technologies, including but not limited to, protein structure information, gene neighbors, sequence composition information, gene expression, physicochemical information, position information, and evolutionary information (Deng, et al., 2014; Guo, Yu, Wen & Li, 2008; Yu, et al., 2016). Zhang et al. (Zhang & Tang, 2016) proposed a PPI prediction method based on gene ontology. However, when structure information cannot be in hand, the domain-based method does not work. Kovács et al. (Kovács, et al., 2019) used network paths of length three to perform link prediction. This approach can offer structural and evolutionary reference to detect protein-protein interactions. Lian et al. (Lian, Yang, Li, Fu & Zhang, 2019) proposed a machine-learning-based predictor for human-bacteria PPIs. This approach introduced two network-property-related feature extraction methods. Then, individual random forest model was constructed for each feature encoding scheme. Finally, the noisy-OR algorithm was employed to predict human-bacteria PPIs. The results on benchmark datasets reveal that the introduced NetTP and NetSS encoding methods could represent important network topology properties. Zahiri et al. (Zahiri, Yaghoubi, Mohammad-Noori, Ebrahimpour & Masoudi-Nejad, 2013) extracted evolutionary information via PPIevo from the position-specific scoring matrix (PSSM) and received better performance and robustness on the HPRD dataset. Hamp et al. (Hamp & Rost, 2015) inferred PPIs based on evolutionary information and SVM.

To improve the PPIs prediction, it is necessary to integrate multiple features mentioned above. Zhang et al. (Zhang, Yu, Xia & Wang, 2019) integrated different descriptors to obtain complimentary information. The constructed ensemble predictor was valid for interactions prediction. Yadav et al. (Yadav, Ekbal, Saha, Kumar & Bhattacharyya, 2019) constructed Bi-LSTM model based on stacked algorithm for the identification of PPIs, which combined multiple levels features using shortest dependency path. Then the information via embedding layer were input into the stacked Bi-LSTM model. The dimensional reduction methods were also performed for effective feature selection, and prediction accuracy improvement since too many features usually bring in additional noise and increase the time complexity in practical problems. You et al. (You, et al., 2014) utilized multi-scale continuous and discrete and minimum redundancy maximum relevance (mRMR) to characterize PPIs coding information. Evaluation indicates mRMR did enhance the success of PPIs prediction and reduce the computation complexity.

Recently, Hashemifar et al. (Hashemifar, Neyshabur, Khan & Xu, 2018) proposed sequence-based convolutional neural networks learning to infer PPIs called DPPI, and deep learning (DL) obtained the high-level and essential feature representations from PSSM. Lei et al. (Lei, et al., 2019) presented a multimodal deep polynomial network called MDPN. For the first stage, high-level features were produced using deep polynomial network based on BLOSUM62, hydrophobic. For the second stage, extreme leaning machine was to predict PPIs through layer-by-layer training. Chen et al (Chen, et al., 2019) presented a PPIs predictive framework PIPR using siamese residual RCNN. This architecture can extract local and contextualized information. However, DL also has the following limitations: (*i*) the number of layers and the number of nodes of the neural network need to be determined before training the DL model (Krizhevsky, Sutskever & Hinton, 2012); (*ii*) the to-be-optimized parameters of DL are diverse on different data, requiring substantial efforts in adjusting the parameters (Krizhevsky, Sutskever & Hinton, 2012; Lecun, Bottou, Bengio & Haffner, 1998; Simonyan & Zisserman, 2015); and (*iii*) DL requires a lot of data for training (Silver, et al., 2018).

Tree ensemble methods have good properties and achieve excellent performance. For example, Feng et al. (Feng & Zhou, 2017) proposed a tree ensemble AutoEncoder (eforest), which can do backward reconstruction using tree-based approach (maximal compatible rule). They utilized forest to perform the process of encoding and decoding for the first time. The experimental results showed that eforest can effectively eliminate noisy information compared with the autoencoder network. Feng et al. (Feng, Yu & Zhou, 2018) proposed a multi-layered GBDT (mGBDT), which can effectively learn hierarchical features through stacking multiple layers. The deep forest (DF) model had fewer hyper-parameters setting and higher flexibility than DL (Zhou & Feng, 2017; Zhou & Feng, 2018). It can deal with non-differential issues without requiring backpropagation algorithms and learn high-level feature information through cascade structure to avoid overfitting. The cascade structure of DF can extract high-level feature information from raw PPIs feature space, and the probability output of upper level with raw features are used as the input of the next level. Specifically, the multi-grained cascade forest is great and robust, hence, can be effectively used to handle machine learning problems such as classification in PPI prediction.

We propose a new PPI prediction method based on DF, so-called GcForest-PPI, where <u>GcForest</u> represents multi-<u>G</u>rained <u>C</u>ascade <u>Forest</u>. The physicochemical information, sequence information, and evolutionary information are retrieved by PAAC, Auto, MMI, CTD, AAC-PSSM, and DPC-PSSM. What is more, elastic net is used to select variables highly relevant to the category labels and GcForest is implemented to identify PPIs based on the known PPIs. Finally, the five-fold cross-validation shows that GcForest-PPI achieves higher accuracy than the state-of-the-art predictors. Cross-species prediction is performed using *Caenorhabditis elegans, Escherichia coli, Homo sapiens,* and *Mus musculus* as independent datasets with the accuracy of 98.58%, 99.04%, 96.01%, and 96.30%, respectively. We also found that (*i*) the PPIs of a CD9-core network are all predicted successfully; (*ii*) GcForest-PPI can predict PPIs in a crossover network and can reveal the biological functions for the Wnt-related pathway; and (*iii*) the PPIs of the cancer-specific network are also all predicted successfully, providing new ideas for studying the associations of drug-disease and drug-target for developing new drugs of cancer treatment.

## 2. Materials and methods

### 2.1. Datasets

Nine PPIs benchmark datasets are utilized to test GcForest-PPI model. The first set was *S. cerevisiae* from DIP core database (Xenarios, et al., 2002). And all protein pairs were identified by the tool CD-HIT (Li, Jaroszewski & Godzik, 2001). The protein sequences with ≤ 50 residues were removed, and sequence similarity ≥ 40% were filtered. So golden standard positive (GSP) set includes 5,594 protein pairs, which have been tested for reliability by the expression profile reliability (EPR) and paralogous verification method (PVM) (Deane, Salwinski, Xenarios & Eisenberg, 2002). A total of 5,594 protein pairs with different subcellular location were selected as golden standard negative (GSN).

The *H. pylori* dataset was validated using the yeast two-hybrid technique (Rain, et al., 2001) and built up by Martin et al. (Martin, Roe & Faulon, 2005), where 1,458 interacting pairs were set as GSP, and 1,458 non-interacting pairs were set as GSN. *Caenorhabditis elegans* (4,013 interacting protein pairs), *Escherichia coli* (6,954 interacting protein pairs), *Homo sapiens* (1,412 interacting protein pairs) and *Mus musculus* (313 interacting protein pairs) were employed as PPIs independent datasets (Zhou, Gao & Zheng, 2011). A one-core network (16 interacting protein pairs) (Yang, et al., 2006), a Wnt-related pathway crossover network (96 interacting protein pairs) (Stelzl, et al., 2005), and cancer-specific network dataset (108 interacting protein pairs) (Amar, Hait, Izraeli & Shamir, 2015) were adopted to predict PPIs networks based on GcForest-PPI.

### 2.2. Feature extraction

The protein structure can be predicted based on sequence, and then to predict its function. Hence, it is feasible that PPIs can be predicted using sequence-based methods via machine learning. We use six feature coding schemes to obtain the physicochemical information, sequence information and evolutionary information, including pseudo amino acid composition (PAAC), autocorrelation descriptor (Auto), multivariate mutual information (MMI), composition-transition-distribution (CTD), amino acid composition PSSM (AAC-PSSM) and dipeptide composition PSSM (DPC-PSSM).

#### 2.2.1. Physicochemical information

Pseudo-amino acid composition (PAAC) and autocorrelation descriptor (Auto) are utilized to extract the physicochemical and composition information. At present, PAAC has shown good properties in proteomics field (Cui, et al., 2019; Qiu, et al., 2018; Yu, et al., 2017a; Yu, et al., 2017b; Yu, et al., 2018; Yu, et al., 2017c). Auto includes Morean-Broto, Moran, and Geary (Chen, Zhang, Ma & Yu, 2019; Chen, et al., 2018). It represents the physicochemical, position information, and the seven physicochemical properties in Auto can be obtained in Supplementary Table S1. The PAAC encoding feature vector *x_u_* can be defined as:

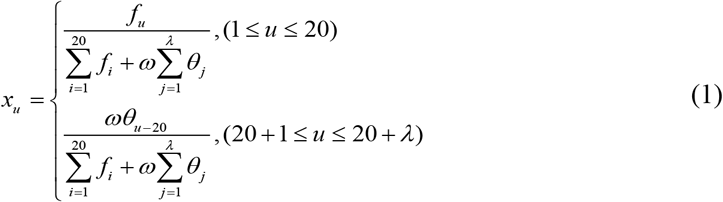

where *f_i_* represents amino acid composition information, *θ_j_* represents layer sequence correlation factor calculated using hydrophobicity, hydrophilicity, and side-chain mass, *ω*=0.05 (Chou, 2001). The shortest length of protein in benchmark PPIs dataset is 12. So the *λ* must satisfy *λ* < 12 and the dimension of PAAC is 20 + *λ*.

We use *A_i_* to characterize the *i-th* amino acids and *P*(*A_i_*) represents the normalized physicochemical values. The 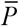 can be employed as the mean value for specific physicochemical property in whole protein sequence. The equation (2), (3), (4) represent Moreau-Broto, Moran, Geary, respectively.

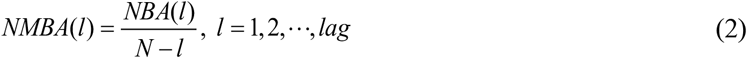

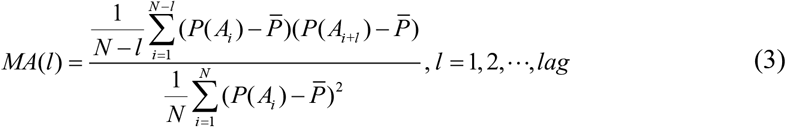

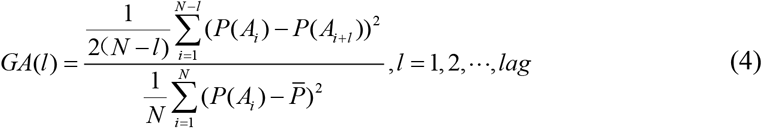

where 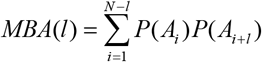, *lag* is the parameter that needs to be adjusted. The dimension of Auto is 3×7×*lag*.

#### 2.2.2. Sequence information

Multivariate mutual information (MMI) (Ding, Tang & Guo, 2016; Ding, Tang & Guo, 2017) and composition-transition-distribution (CTD) are utilized to obtain sequence information (Zhang, Yu, Xia & Wang, 2019). MMI can represent the information entropy and group features. CTD can obtain the distribution pattern and effective sequence information. The groups of amino acids are listed in Supplementary Table S2.

For MMI, the amino acid residues can be classified into seven classes according to Supplementary Table S3. The algorithm flowchart of MMI is shown in the Supplementary Fig. S1. For a given protein sequence, we can define various 2-gram *I*(*a, b*) and 3-gram *I*(*a,b,c*) features. Take “*C*_0_*C*_0_*C*_0_”,“*C*_0_*C*_0_*C*_1_”,⋯,“*C*_6_*C*_6_*C*_6_” for example. The information entropy can be expressed as:

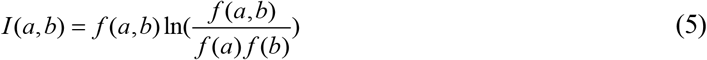

where *f*(*a,b*) represents frequency 2-gram (a, b) for given sequence. *f*(*a*) represents frequency of *a*.

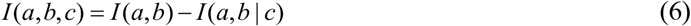

where *a,b,c* are types of amino acid in triplet, and *I*(*a, b*|*c*)=*H*(*a*|*c*)-*H*(*a*|*b,c*) which could be described as:

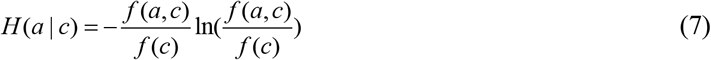

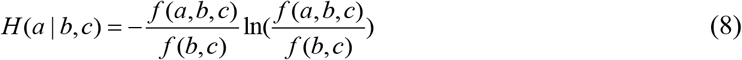

Finally, each protein sequence yields 84-dimensional 3-gram features and 28-dimensional 2-gram features. The dimension of MMI is 119.

In CTD (Chen, et al., 2018), amino acids are grouped into three groups based on hydrophobicity: polar (P), neutral (N), and hydrophobic (H). Using *N*(*r*) represents the character type *r* in the replaced sequence, and *N* is sequence length. Given sequence MTTTVPKVFAFHEF. It can be represented as ‘32223213323213’ according to Hydrophobicity_PRAM900101. ‘1’ represents polar, ‘2’ represents neutral, ‘3’ represents hydrophobicity.

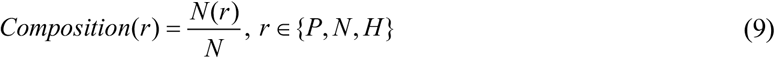

The composition generate grouped information, the frequency of ‘1’ is 2/14 = 0.1429, the frequency of ‘2’ is 6/14 = 0.4286, the frequency of ‘3’ is 6/14 = 0.4286.

The T descriptor first converts the original sequence into a replacement sequence, and T includes three characteristics, the dipeptide composition frequency from the polar group to the neutral group and the composition frequency from the neutral group to the polar group. Transitions between the neutral group and the hydrophobicity and these between hydrophobic group and the polar group are defined in the same way.

The T descriptor is defined as follows:

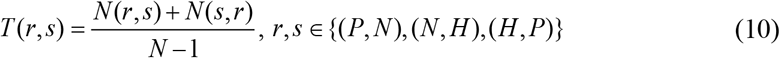

where *N*(*r,s*) represents dipeptide frequency, the value of (*P,N*) is 2/13 = 0.1538, the value of (*N,H*) is 6/13 = 0.4615, the value of (*H,P*) is 2/13 = 0.1538.

For each group (P, N and H), we obtain the pattern information of the first, 25%, 50%, 75% and 100% of the encoded grouped sequence. Take ‘3’ for example, there are 6 residues encoded ‘3’. The first ‘3’ is 1. The second ‘3’ is 25%×6 = 1. The third ‘3’ is 50%× 6 = 3. The fourth ‘3’ is 75% × 6 = 4. The fifth '3 is 100% × 6 = 6. The position in the first, the second, the third, the fourth, the fifth ‘3’ of whole sequence are 1, 1, 8, 9, 14, respectively. So the distribution descriptor for ‘3’ are (1/14), (1/14), (8/14), (9/14), (14/14).

The Composition generates a 39-dimensional sample numeric vector, the Transition generates a 39-dimensional sample numeric vector, and the Distribution generates a 195-dimensional sample numeric vector. In summary, the CTD generates a 273-dimensional sample numeric vector.

#### 2.2.3. Evolutionary information

Evolutionary information in the position-specific scoring matrix (PSSM) is essential in proteomics (Supplementary File S1). The amino acid composition PSSM (AAC-PSSM) and dipeptide composition PSSM (DPC-PSSM) are utilized to generate evolutionary information. Some researchers have used PSSM to leverage encoding information, including the identification of drug-target interaction (Shi, et al., 2019), detecting protein-protein interaction site (Wang, et al., 2019; Wei, Han, Yang, Shen & Yu, 2016; Zhang, Li, Quan, Chen & Q. Lü, 2019).

PSSM are converted to feature vector by AAC-PSSM via equation (11)

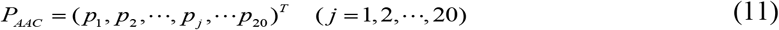

where 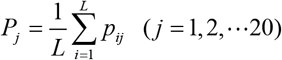, *p_j_* represents the composition evolutionary information of the *j* amino acid residue. And the dimension of AAC-PSSM is 20.

ACC-PSSM only represents the composition information from PSSM, and loses the position information, which is insufficient to fully represent the evolutionary information. DPC-PSSM can reflect the sequence-order information of PSSM, the encoding feature vector can be expressed as

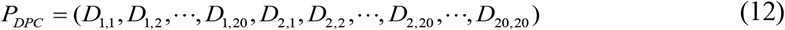

where 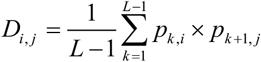, the dimension of DPC-PSSM is 400.

### 2.3. Elastic net

Elastic Net (EN) (Zou & Hastie, 2005) is a feature selection method based on regularization term. EN not only keeps the sparse model of LASSO but also maintains the regularization properties of the ridge regression. *α, β* are penalty terms, which represent a compromise strategy between LASSO and ridge regression. The objective function of the elastic net is

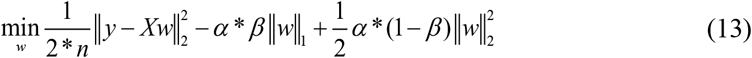

where *X* is the sample matrix, *y* is the category label, *n* represents sample number, and *w* indicates the regression coefficient. The L1 regular term is used to generate the sparse model (LASSO), and the L2 regularization can produce a group effect.

### 2.4. Deep forest

Deep forest (DF) is a forest-based ensemble learning method for trees (Zhou & Feng, 2017; Zhou & Feng, 2018), which can represent high-level feature information by cascade structure. Zhou et al. used two random forest (RF) (Breiman, 2001), and two extremely randomized trees (Extra-Trees) (Geurts, Ernst & Wehenkel, 2006) to construct the deep forest. Considering the boosting algorithm achieves higher computation accuracy and better model generation ability. Especially the XGBoost (Chen & Guestrin, 2016), which combines the linear model, regularized objective and second-order approximation via boosting algorithm to avoid over-fitting, reduce computational costs, enhance predictive performance. Meanwhile, sub-samples speed up the parallel computing in the process of tree learning. So we develop a new deep forest architecture to implement GcForest, which is composed of four XGBoost, four RF and four Extra-Trees. XGBoost is a variant of gradient boosting decision tree whose base classifier is regression tree. The base classifiers of the RF and Extra-Trees are decision tree. In this way, an outstanding deep forest contain good and diverse base classifier. Then the deep layer architecture GcForest can obtain complementary advantages and essential features.

XGBoost is an ensemble algorithm. Given dataset 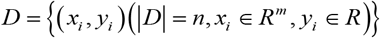, the loss function of XGBoost is shown as

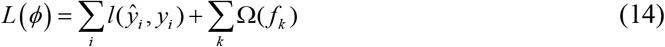

where *L* is the convex objective function, Ω penalizes the complexity of XGBoost, *f_k_* represents the *k-th* regression tree. Then, second-order Taylor is adopted to enhance the predictive performance:

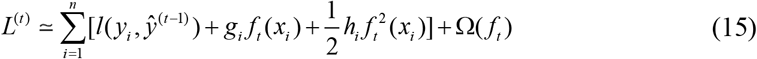

where 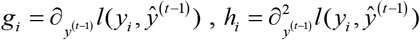 represents the first order and second order gradient statistics.

RF is a bagging ensemble classifier using random bootstrap. Gini coefficient is employed as the evaluation to split the node for tree learning. There are two main differences between Extra-Trees and RF. (*i*) Extra-Trees uses all training set to generate decision tree. (*ii*) Each tree is segmented and grown at each node by randomly selecting a feature.

The levels include four XGBoost, four RF, and four Extra-Trees. The cascade structure is shown in Fig. 1A. Suppose there are two classes to predict, each forest will output a two-dimensional class vector, and each layer will generate a 24-dimensional new class vector. The newly generated class vectors are concatenated with the raw protein feature vectors to produce multi-level features. The output class probability score of the last layer is shown in Fig. 1B.

**Fig. 1.**
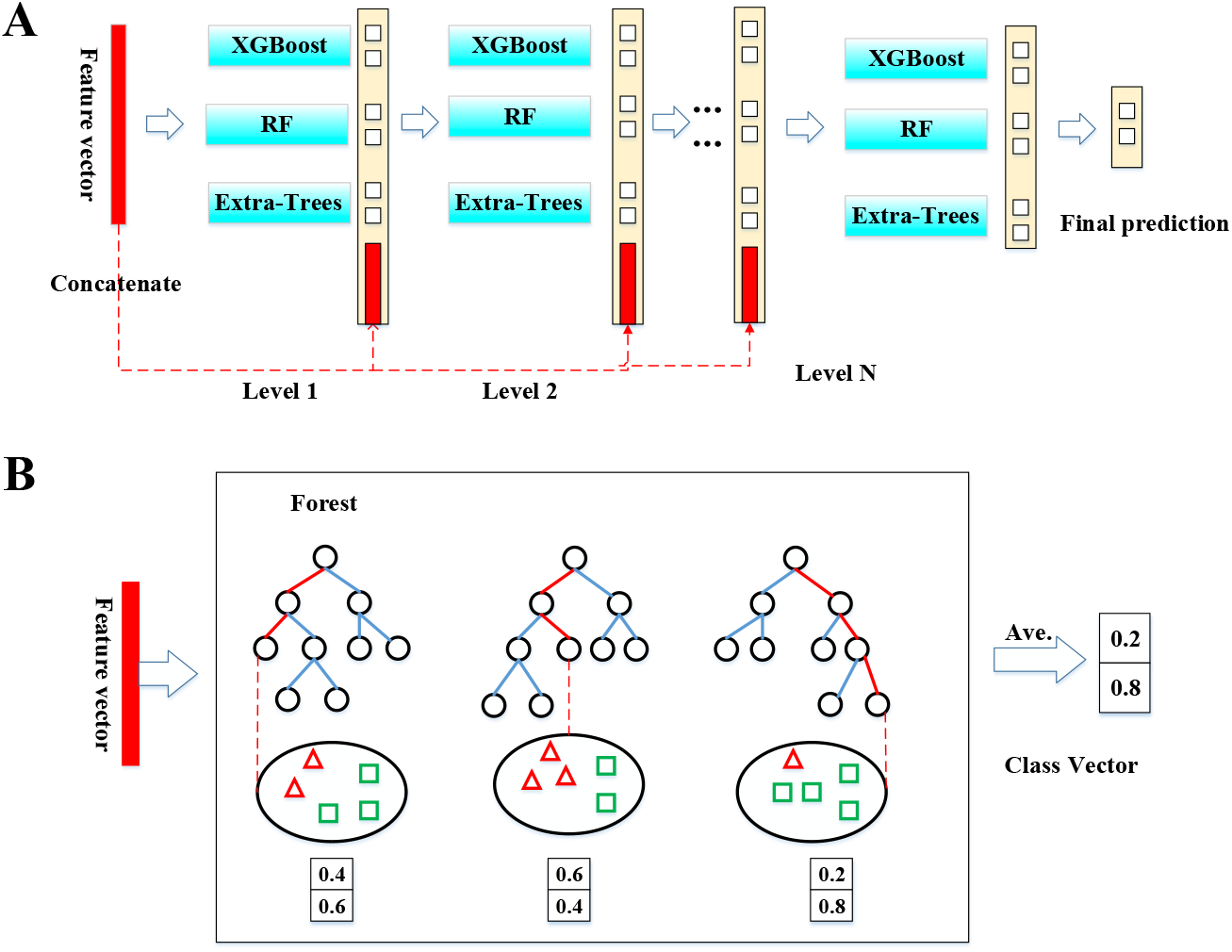
The GcForest structure and the generation of a class vector. (A) Illustration of GcForest structure. (B) The generation of class vector. Different marks in leaf nodes represent different classes.

As illustrated in Fig. 1, given an instance, each forest can produce an estimate of class distribution by calculating the percentage of different types of training samples at the leaf node. The number of iterations on XGBoost is set to 500. The RF includes 500 decision trees and randomly selects 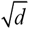 features as candidate subsets (*d* is the dimension of dataset). The Extra-Trees consist of 500 trees.

To reduce overfitting of GcForest, the class vector generated by each forest using five-fold cross-validation. Specifically, each sample will be employed as training set twelve times. Then, the class vectors are concatenated to produce augmented class vectors. The feature information is obtained from known sequences in the previous study, but they may generate noisy data inputs. It is reasonable to extract high-level feature information for prediction, and the probability output is employed as the next level of the forest. So, DF has good generalization ability, and the deep structure can exploit potential information from PPIs.

### 2.5. Performance evaluation and model construction

In order to evaluate the GcForest-PPI model, the evaluation indicators included Recall, Precision, Accuracy (ACC) and Matthews correlation coefficient (MCC) (Cui, et al., 2019; Du, et al., 2017; Tian, et al., 2019; Yu, et al., 2018).

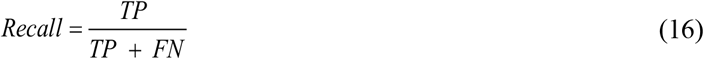

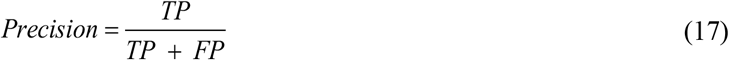

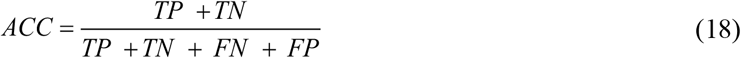

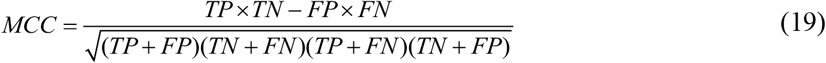

where TP, TN, FP, and FN represent true positive, true negative, false positive, and false negative, respectively. Receiver operating characteristic (ROC) curve (Shi, et al., 2019; Wang, et al., 2019) and AUC, Precision recall curve (PR) (Davis & Goadrich, 2006) and AUPR are also indicators to assess the predictive performance of GcForest-PPI. The workflow of GcForest-PPI is shown in Fig. 2 with detailed steps described as follows.

***Step 1:*** Protein pairs. We collect six PPIs dataset. Input interacting pairs and non-interacting pairs.
***Step 2:*** Feature extraction. The effective initial coding information of PPIs could be obtained by PAAC, Auto, MMI, CTD, AAC-PSSM and DPC-PSSM. These descriptors can produce complimentary information by integrating physicochemical, position, sequence, composition and evolutionary information.
***Step 3:*** Feature selection. The elastic net based on L1 and L2 regularization can eliminate redundancy and retain essential variables. Adjusting the parameters *α* and *β* via five-fold cross-validation to generate effective subset for identifying PPIs. The comparison indicates elastic net obviously outperforms other dimensional reduction approaches.
***Step 4:*** Deep forest and model construction. The important feature representations can be obtained for binary PPIs prediction task via Step 2 and Step 3. Then ensemble XGBoost, RF and Extra-Trees via cascade architecture to implement the task, and the predictive tool GcForest-PPI for PPIs based on deep forest is built up.
***Step 5:*** Model evaluation. We apply GcForest-PPI on four cross-species datasets, CD9-core network, crossover network and cancer-specific network. Then list the comparison of GcForest-PPI with the state-of-the-art predictors and plot the three types of protein-protein interactions networks.

**Fig. 2.**
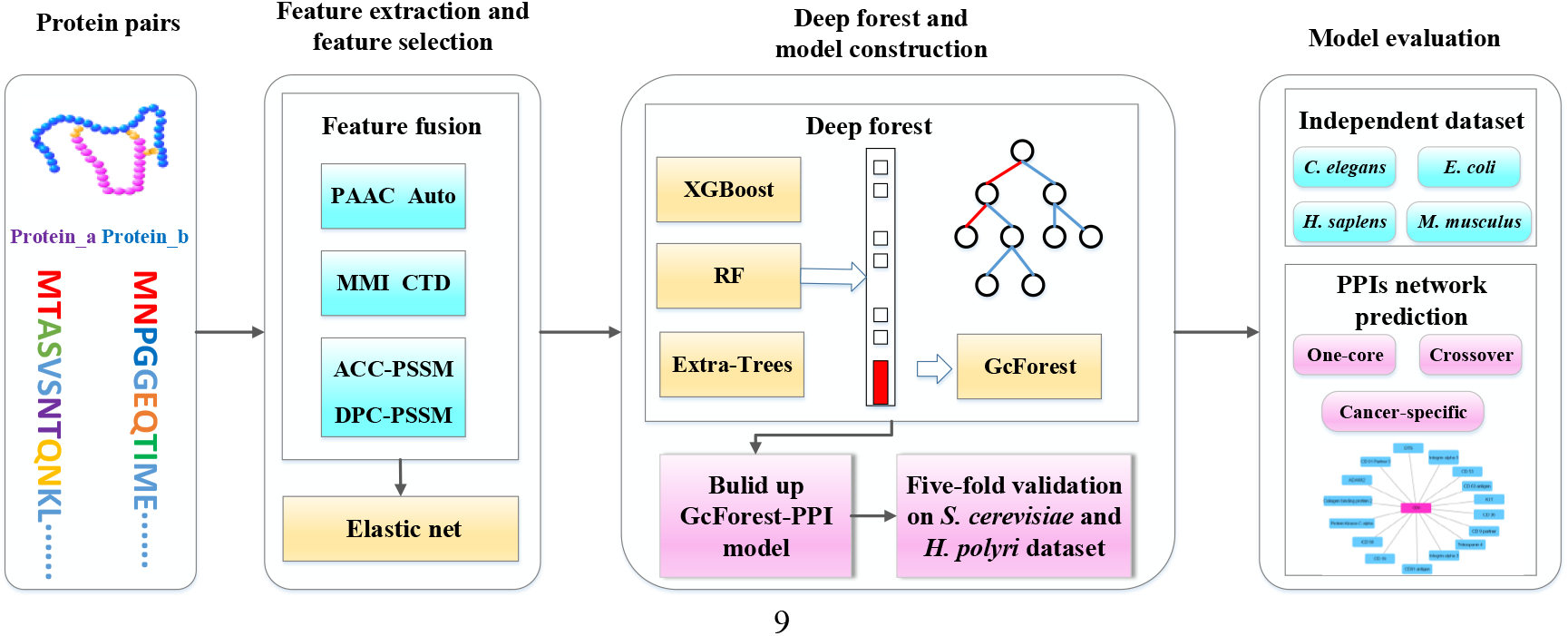
The overall framework of GcForest-PPI. First, input the protein pairs and utilize PAAC, Auto, MMI, CTD, AAC-PSSM and DPC-PSSM to encode feature values. Then using elastic net to find effective, significant, and valuable feature subset. Finally, the GcForest-PPI model is constructed based on deep forest. The output of GcForest-PPI should decide whether protein pairs are PPIs or non-PPIs.

## 3. Results and discussion

All simulation results of GcForest-PPI were performed on Windows Server 2012R2 with 32.0GB of RAM, GcForest-PPI was implemented by Python 3.6 and MATLAB.

### 3.1. Parameter selection of the feature extraction

The parameter *λ* in PAAC indicates the order information in the coding process. The parameter *lag* represents the interval of two residues in the computational process of AD. For different *λ* and *lag* values, deep forest is adopted to construct the predictor. The prediction results are listed in Supplementary Table S4 and Supplementary Table S5. The intuitive parameter changes of accuracy are shown in Fig. 3.

**Fig. 3.**
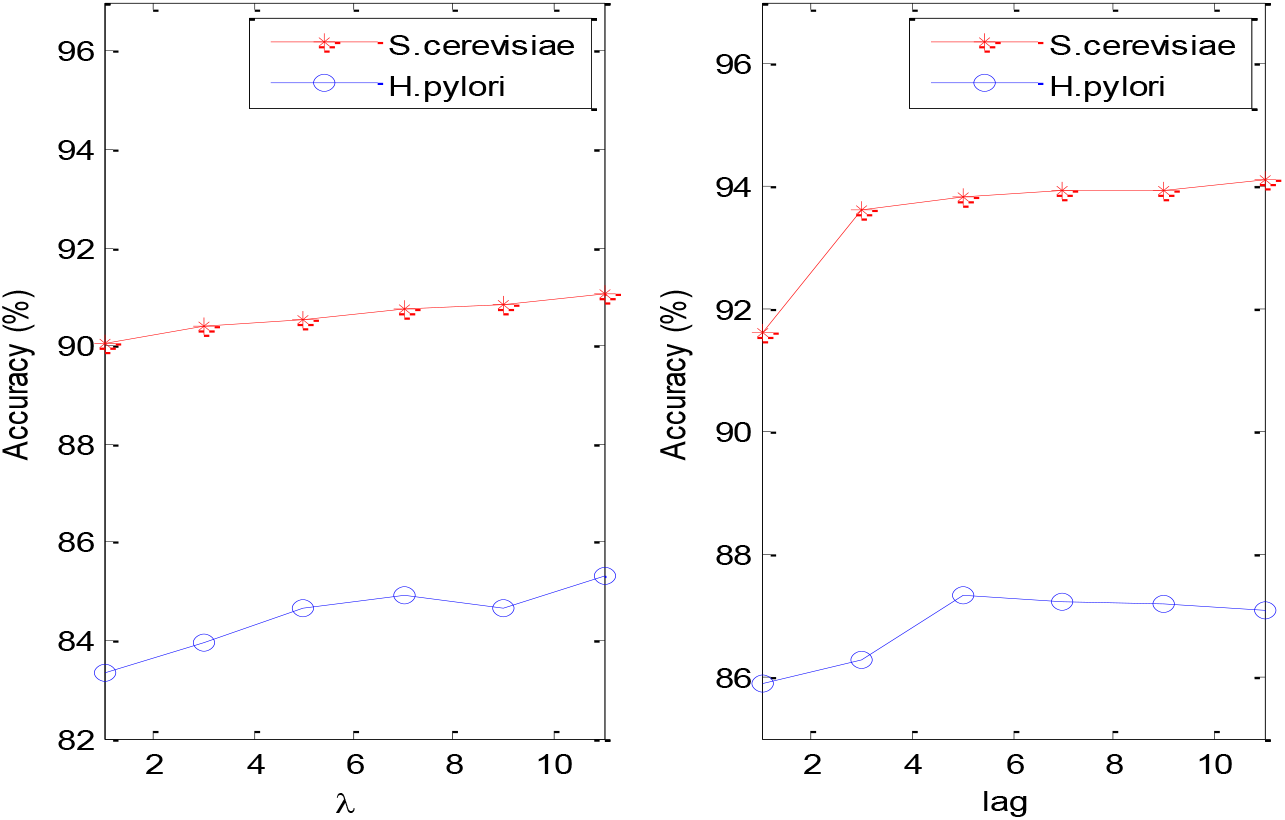
The parameter optimization of PAAC and Auto for *S. cerevisiae* and *H. pylori*. The *λ* represents the parameter need to be adjusted in PAAC. The *lag* represents the parameter need to be adjusted in AD.

As shown in Fig. 3, we can notice that the changes of *λ* and *lag* can effect the prediction condition. For the PAAC, the peaks of the *S. cerevisiae* and *H. pylori* datasets are same. Hence, we determine *λ* = 11 in PAAC. For the Auto algorithm, the peak point of *S. cerevisiae* is 11, and the peak point of *H. pylori* is 5. Considering that we use the *S. cerevisiae* dataset as the train set to predict the independent test set, we set *lag*=11 to unify the parameter *lag*. PAAC and Auto can mine the sequence physicochemical information. MMI and CTD obtain sequence and composition pattern through grouping amino acids. The PSSM can be converted to important evolutionary representation related to PPIs through AAC-PSSM and DPC-PSSM. For each protein sequence, six feature coding schemes are combined to obtain 1,074 features. Then protein pair vectors are concatenated to fully characterize pairwise relations whose dimension is 2148.

### 3.2. Elastic net performs better than other dimensionality reduction method

The elastic net feature selection was employed to optimize the feature set. From Supplementary File S2, we can see the parameters of the elastic net are *α*=0.03 and *β*=0.1. The numbers of optimal features are 476 and 516 on *S. cerevisiae* and *H. pylori*, respectively. And the raw features and optimal features from different feature information are shown in Supplementary Fig. S2. What is more, we also use principal component analysis (PCA) (Wold, Esbensen & Geladi, 1987), kernel principal component analysis (KPCA) (Schölkopf, Smola & Müller, 1998), local linear embedding (LLE) (Roweis & Saul, 2000), spectral embedding (SE) (Ng, Jordan & Weiss, 2002), singular value decomposition (SVD) (Wall, Rechtsteiner & Rocha, 2002), semi-supervised dimensionality reduction (SSDR) (Zhang, Zhou & Chen, 2007) to eliminate redundant information. Then construct the GcForest-PPI framework based on deep forest via five-fold cross-validation. The main experimental results of *S. cerevisiae*, and *H. pylori* are shown in Supplementary Table S8. The ROC curves, PR curves and AUC values, AUPR values are shown in Fig. 4. and Supplementary Table S9, respectively.

**Fig. 4.**
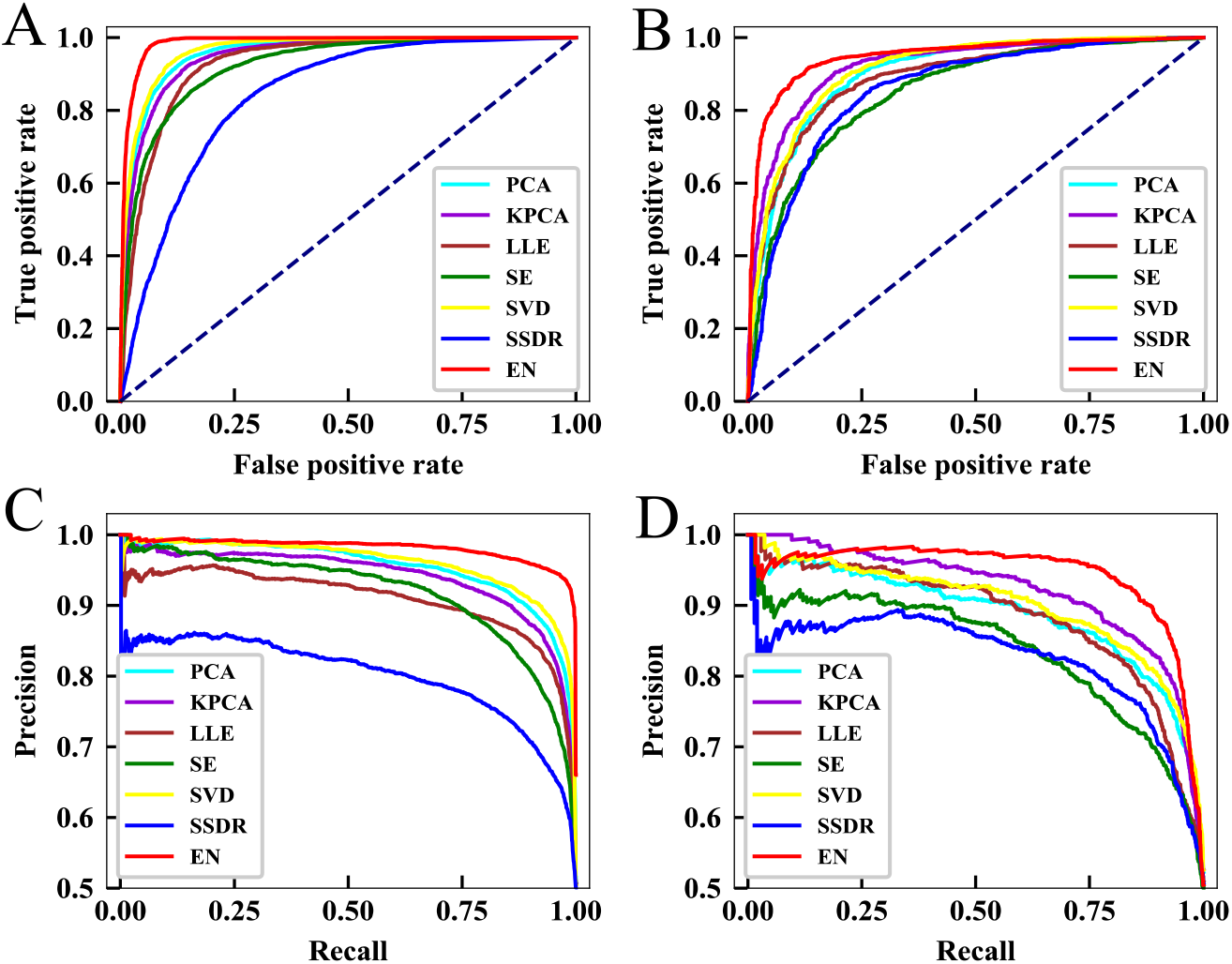
Predictive performance of PCA, KPCA, LLE, SE, SVD, SSDR and EN via five-fold cross-validation. (A-B) The ROC curves of *S. cerevisiae* and *H. pylori*. (C-D) The PR curves of *S. cerevisiae* and *H. pylori*.

It is noticed that from Fig. 4A, the accuracy of EN exceeds the PCA, KPCA, LLE, SE, SVD, SSDR (0.9864 vs. 0.9603, 0.9497, 0.9302, 0.9243, 0.9664, 0.8425) for *S. cerevisiae*. EN is 2.61% higher than PCA (0.9864 vs. 0.9603) and 14.39% higher than SSDR (0.9864 vs. 0.8425). As Fig. 4B shows compared with other methods, the robustness of the elastic net is optimal. The AUC value of EN outperforms PCA, KPCA, LLE, SE, SVD, SSDR (0.9816 vs. 0.9545, 0.9402, 0.9088, 0.9172, 0.9605, 0.7999). From the PR curve of Fig. 4C, EN achieves relatively high accuracy compared with PCA, KPCA, LLE, SE, SVD, SSDR (0.9485 vs. 0.9019, 0.9230, 0.8888, 0.8509, 0.9129, 0.8560) in terms of AUPR. Fig. 4D plots the PR curve and the AUPR of EN is 2.4%-11.95% higher than PCA, KPCA, LLE, SE, SVD, SSDR (0.9449 vs. 0.8873, 0.9209, 0.8845, 0.8356, 0.9007, 0.8254) on the *H. pylori*. And the AUPR of EN is 5.76% higher than PCA (0.9449 vs. 0.8873). Therefore, EN can effectively eliminate redundant information that has little correlation with PPIs, and retain important feature subset information, which provides effective feature fusion information for deep forest.

### 3.3. GcForest performs better than other classifiers

To verify the effectiveness of DF, we also use logistic regression (LR) (Yu, Huang & Lin, 2011), Naïve Bayes (NB) (Friedman, Geiger & Pazzanzi, 1997), K nearest neighbors (KNN) (Nigsch, et al., 2006), AdaBoost (Freund & Schapire, 1997), random forest (RF) (Breiman, 2001) and SVM (Cortes & Vapnik, 1995) six classifiers to predict PPIs. The main prediction results of the five-fold cross-validation on the *S. cerevisiae* and *H. pylori* datasets are shown in Supplementary Table S10. The results of the ROC curves and the PR curves, AUC and AUPR are shown in Fig. 5, Table S11, respectively.

**Fig. 5.**
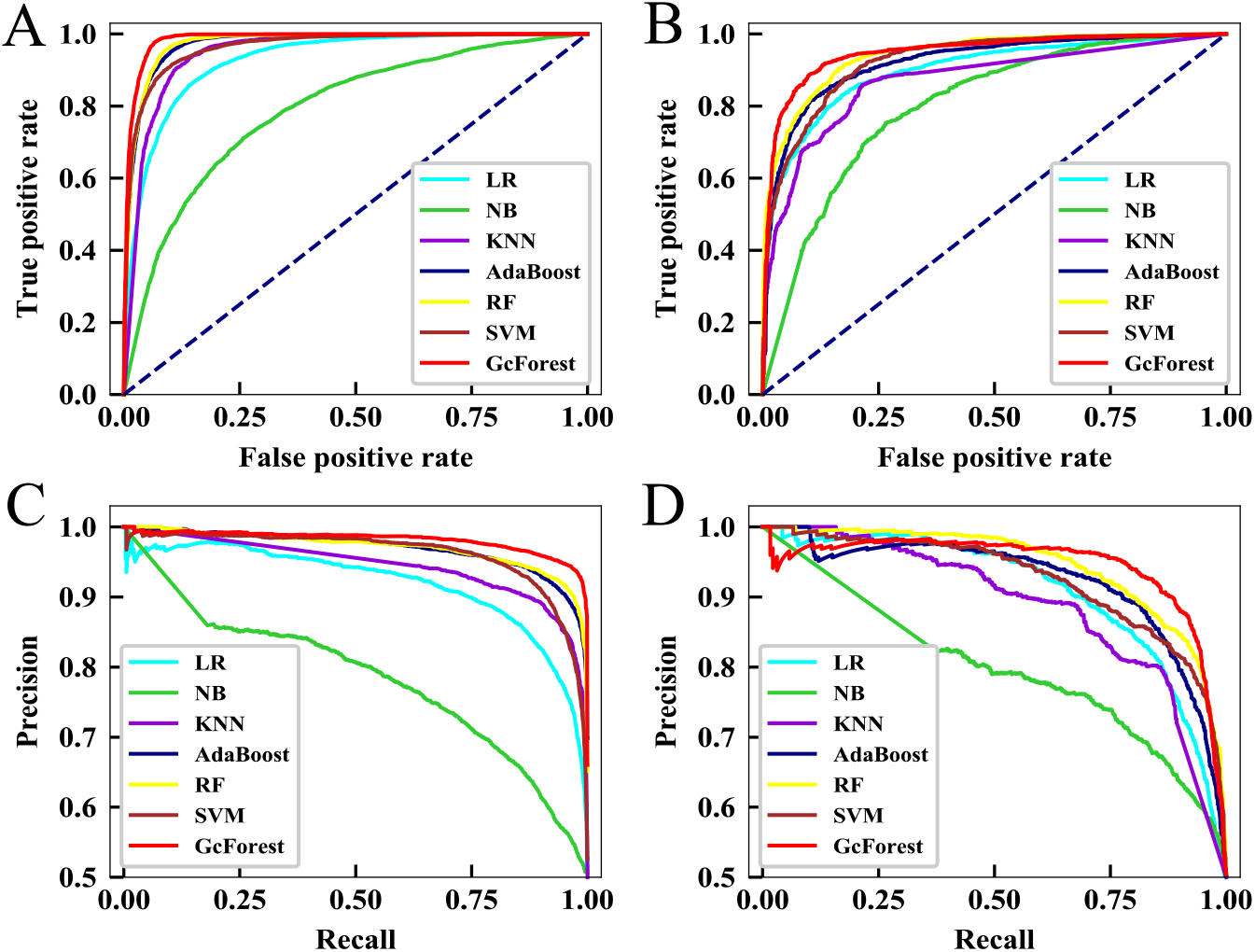
Predictive performance of LR, NB, KNN, AdaBoost, RF, SVM, and GcForest via five-fold cross-validation. (A-B) The ROC curves illustrating the prediction of *S. cerevisiae* and *H. pylori*. (C-D) The PR curves representing the performance on the *S. cerevisiae* and *H. pylori*.

From Fig. 5A, the ROC curve of DF performs the best on *S. cerevisiae* compared with LR, NB, KNN, AdaBoost, RF, SVM classifiers (0.9864 vs. 0.9298, 0.7914, 0.9503, 0.9750, 0.9762, 0.9653). The AUC of GcForest is increased by 2.11% over SVM (0.9864 vs. 0.9653). From Fig. 5B, the AUC value of GcForest is higher than LR, NB, KNN, AdaBoost, RF, SVM (0.9816 vs. 0.9189, 0.7721, 0.9270, 0.9693, 0.9704, 0.9616). GcForest is 1.12%-20.95% higher than the other six machine-learning-based algorithms. From Fig. 5C, the PR curve indicates that GcForest is superior to LR, NB, KNN, AdaBoost, RF, SVM for predicting PPIs (0.9485 vs. 0.8996, 0.8022, 0.8722, 0.9225, 0.9427, 0.9264) in terms of AUPR on *S. cerevisiae*. GcForest is 0.58%-14.63% higher than the other six classifiers. Fig. 5D indicates the AUPR of GcForest is higher than LR, NB, KNN, AdaBoost, RF, SVM (0.9449 vs. 0.9091, 0.7653, 0.8764, 0.9229, 0.9447, 0.9246). GcForest is 3.58%, 2.03% higher than LR, SVM (0.9449 vs. 0.9091), (0.9449 vs. 0.9246).

We use DF to predict PPIs using XGBoost, random forest and extremely randomized trees to construct cascade forest for the first time. The high-level feature information can be extracted, and probability output from the previous layer is input into the next level. The experimental results show that GcForest is superior to LR, NB, KNN, AdaBoost, RF and SVM classifiers. Tree-ensemble methods can mine the potential feature information of protein interaction pairs through layer-by-layer learning, and thus it can fit the non-linear relationship to determine whether a pair is interacting or non-interacting. DF can have flexible hyperparameter adjustment, high efficiency, and good scalability.

### 3.4. Comparison with other state-of-the-art PPIs prediction methods

To verify the validity of the GcForest-PPI model, we listed the results of ACC+SVM (Guo, Yu, Wen & Li, 2008), Code4+KNN (Yang, Xia & Gui, 2010), LD+SVM (Zhou, Gao & Zheng, 2011), MCD+SVM (You, et al., 2014), LRA+RF (You, Li & Chan, 2017), DeepPPI (Du, et al., 2017), DPPI (Hashemifar, Neyshabur, Khan & Xu, 2018) on *S. cerevisiae* in Table 1, and the results of SVM (Martin, Roe & Faulon, 2005), Ensemble of HKNN (Nanni & Lumini, 2006), DCT+WSRC (Huang, You, Xin, Leon & Wang, 2015), MCD+SVM (You, et al., 2014), MIMI+ NMBAC+RF (Ding, Tang & Guo, 2016), PCA-EELM (You, Lei, Zhu, Xia & Wang, 2013), DeepPPI (Du, et al., 2017) on *H. pylori* in Table 2.

**Table 1.** Comparison of different PPIs prediction methods on *S. cerevisiae* dataset.

| Method | ACC (%) | Recall (%) | Precision (%) | MCC |
| --- | --- | --- | --- | --- |
| ACC+SVM <sup>a</sup> | 89.33 ± 2.67 | 89.93 ± 3.68 | 88.87 ± 6.16 | N/A |
| Code4+KNN <sup>b</sup> | 86.15 ± 1.17 | 81.03 ± 1.74 | 90.24 ± 1.34 | N/A |
| LD+SVM <sup>c</sup> | 88.56 ± 0.33 | 87.37 ± 0.22 | 89.50 ± 0.60 | 0.7715 ± 0.0068 |
| MCD+SVM <sup>d</sup> | 91.36 ± 0.36 | 90.67 ± 0.69 | 91.94 ± 0.62 | 0.8421 ± 0.0059 |
| LRA+RF <sup>e</sup> | 94.14 ± 1.8 | 91.22 ± 1.6 | 97.10 ± 2.1 | 0.8896 ± 0.026 |
| DeepPPI <sup>f</sup> | 94.43 ± 0.30 | 92.06 ± 0.36 | 96.65 ± 0.59 | 0.8897 ± 0.0062 |
| DPPI <sup>g</sup> | 94.55 | 92.24 | 96.68 | N/A |
| GcForest-PPI | 95.44 ± 0.18 | 92.72 ± 0.44 | 98.05 ± 0.25 | 0.9102 ± 0.0035 |

Note: N/A means not available. The values behind ± represent the standard deviation. <sup>a</sup>
Results reported by (Guo, Yu, Wen & Li, 2008) and the paper uses the holdout validation. <sup>b</sup> Results reported by (Yang, Xia & Gui, 2010). <sup>c</sup> Results reported by (Zhou, Gao & Zheng, 2011). <sup>d</sup> Results reported by (You, et al., 2014). <sup>e</sup> Results reported by (You, Li & Chan, 2017). <sup>f</sup> Results reported by (Du, et al., 2017). <sup>g</sup> Results reported by (Hashemifar, Neyshabur, Khan & Xu, 2018).

From Table 1, we can see the GcForest-PPI model achieves the best prediction performance with an ACC of 95.44%, Recall of 92.72%, Precision of 98.05%, and MCC of 0.9102. The ACC and Recall based on GcForest-PPI, DPPI (Hashemifar, Neyshabur, Khan & Xu, 2018), DeepPPI (Du, et al., 2017) and ACC+SVM (Guo, Yu, Wen & Li, 2008) are (95.44% and 92.72%), (94.55% and 92.24%), (94.43% and 92.06%) and (89.33% and 89.93%). The ACC of GcForest-PPI is 1.01% higher than DeepPPI (Du, et al., 2017) (95.44% vs. 94.43%). On Recall, GcForest-PPI is 1.5% higher than the LRA+RF (You, Li & Chan, 2017) (92.72% vs. 91.22%). On Precision, GcForest-PPI is 9.18% higher than the ACC+SVM (Guo, Yu, Wen & Li, 2008) (98.05% vs. 88.87%). In summary, our proposed method GcForest-PPI is powerful on *S. cerevisiae* for PPIs identification.

From Table 2, we can see that on the *H. pylori*, The ACC and Recall of GcForest-PPI, PCA-EELM (You, Lei, Zhu, Xia & Wang, 2013), MCD+SVM (You, et al., 2014), DCT+WSRC (Huang, You, Xin, Leon & Wang, 2015), are (89.26% and 89.71%), (87.50% and 88.95%), (84.91% and 83.24%) and (86.74% and 86.43%). The ACC of GcForest-PPI is 3.03%, 5.86%, 1.67% higher than DeepPPI (Du, et al., 2017), SVM (Martin, Roe & Faulon, 2005) and MIMI+ NMBAC+RF (Ding, Tang & Guo, 2016), respectively (89.26% vs 86.23%, 83.40%, 87.59%). From Recall, we can see GcForest-PPI is 3.28% higher than DCT+WSRC (Huang, You, Xin, Leon & Wang, 2015) (89.71% vs. 86.43%). The Precision and MCC of GcForest-PPI, DeepPPI (Du, et al., 2017), MMI+NMBAC+RF (Ding, Tang & Guo, 2016) are (88.95% and 0.7857), (84.32% and 0.7263) and (88.23% and 0.7524). On MCC, GcForest-PPI is 0.44%-5.94% higher than other PPIs prediction tools. GcForest-PPI is 5.94% higher than DeepPPI (Du, et al., 2017) (0.7857 vs. 0.7263).

**Table 2.** Comparison of different PPIs prediction methods on *H. pylori* dataset.

| Method | ACC (%) | Recall (%) | Precision (%) | MCC |
| --- | --- | --- | --- | --- |
| SVM <sup>a</sup> | 83.40 | 79.90 | 85.70 | N/A |
| Ensemble of HKNN <sup>b</sup> | 86.60 | 86.70 | 85.00 | N/A |
| DCT+WSRC <sup>c</sup> | 86.74 | 86.43 | 87.01 | 0.7699 |
| MCD+SVM <sup>d</sup> | 84.91 | 83.24 | 86.12 | 0.7440 |
| MIMI+ NMBAC+RF <sup>e</sup> | 87.59 | 86.81 | 88.23 | 0.7524 |
| PCA-EELM <sup>f</sup> | 87.50 | 88.95 | 86.15 | 0.7813 |
| DeepPPI <sup>g</sup> | 86.23 | 89.44 | 84.32 | 0.7263 |
| GcForest-PPI | 89.26±1.07 | 89.71±2.26 | 88.95±1.36 | 0.7857±0.0212 |

*Note:* N/A means not available. The values behind ± represent the standard deviation. <sup>a</sup> Results reported by (Martin, Roe & Faulon, 2005) and this paper uses ten-fold cross-validation. <sup>b</sup> Results reported by (Nanni & Lumini, 2006) and this paper uses ten-fold cross-validation. <sup>c</sup> Results reported by (Huang, You, Xin, Leon & Wang, 2015), and this paper used ten-fold cross-validation. <sup>d</sup> Results reported by (You, et al., 2014). <sup>e</sup> Results reported by (Ding, Tang & Guo, 2016). <sup>f</sup> Results reported by (You, Lei, Zhu, Xia & Wang, 2013). <sup>g</sup> Results reported by (Du, et al., 2017).

### 3.5. Prediction results on four independent species

The pros and cons of GcForest-PPI are further evaluated on *C. elegans* (4,013 interacting protein pairs), *E. coli* (6,954 interacting protein pairs), *H. sapiens* (1,412 interacting protein pairs), and *M. musculus* (313 interacting protein pairs) and the whole samples of the *S. cerevisiae* are regarded as training set. The results of GcForest-PPI and DPPI (Hashemifar, Neyshabur, Khan & Xu, 2018), DeepPPI (Du, et al., 2017), MLD+RF (You, Chan & Hu, 2015), DCT+WSRC (Huang, You, Xin, Leon & Wang, 2015) are shown in Table 3.

**Table 3.** Comparison of performance of the proposed method with other state-of-the-art predictors on the independent dataset.

| Species | ACC (%) |  |  |  |  |
| --- | --- | --- | --- | --- | --- |
|  | GcForest-PPI | DPPI <sup>a</sup> | DeepPPI <sup>b</sup> | MLD+RF <sup>c</sup> | DCT+WSRC <sup>d</sup> |
| *H. sapiens* | 98.58 | 96.24 | 94.84 | 94.19 | 82.22 |
| *M. musculus* | 99.04 | 95.84 | 92.19 | 91.96 | 79.87 |
| *C. elegans* | 96.01 | 95.51 | 93.77 | 87.71 | 81.19 |
| *E. coli* | 96.30 | 96.66 | 91.37 | 89.30 | 66.08 |

Note: <sup>a</sup> Results reported by (Hashemifar, Neyshabur, Khan & Xu, 2018). <sup>b</sup> Results reported by (Du, et al., 2017). <sup>c</sup> Results reported by (You, Chan & Hu, 2015). <sup>d</sup> Results reported by (Huang, You, Xin, Leon & Wang, 2015).

From Table 3, we can know that the ACC of GcForest-PPI on *H. sapiens, M. musculus, C. elegans*, and *E. coli* are 98.58%, 99.04%, 96.01%, and 96.30%, respectively. GcForest-PPI is superior to the DPPI on *H. sapiens, M. musculus* and *C. elegans* (98.58% vs. 96.24%), (99.04% vs. 95.84%), and (96.01% vs. 95.51%). At the same time, the GcForest-PPI performs better than DeepPPI (Du, et al., 2017), MLD+RF (You, Chan & Hu, 2015), and DCT+WSRC (Huang, You, Xin, Leon & Wang, 2015). This shows that GcForest-PPI model characterizes PPIs information using *S. cerevisiae* dataset. In other words, it is possible that PPIs of one species can predict cross-species and the co-evolved relationship can be mined via cascade structure based on XGBoost, RF and Extra-Trees.

### 3.6. PPIs network prediction

We use the one-core network, Wnt-related pathway network and cancer-specific network to evaluate the advantages and disadvantages of the GcForest-PPI model. It provides some reference for identifying PPIs from unknown PPIs networks. The one-core network is a simple CD9-core network including 17 genes. The second is a crossover network for Wnt-related pathway. This network has 78 genes consisting of 96 PPI pairs. The cancer-specific network (Amar, Hait, Izraeli & Shamir, 2015) consists of 78 genes, which are of importance in DNA replication and cancer pathways. The interaction pairs in the cancer-specific network are derived from the IntAct database (Kerrien, et al., 2007).

The GcForest-PPI prediction model is constructed using the *S. cerevisiae* dataset to predict the one-core network with CD9 as the core protein, the Wnt-related pathway network and the cancer-specific network. According to the discussion in Section 3.1, the protein pairs are converted to 2,148-dimensional feature vector by PAAC, Auto, MMI, CTD, AAC-PSSM, and DPC-PSSM (where *λ* is 11 in PAAC and *lag* is 11 in Auto).Then we select 476 important features via elastic net. Finally, deep-forest-based model GcForest-PPI using random forest, Extra-trees and XGBoost is constructed. The results of three types PPIs networks are shown in Fig. 6.

**Fig. 6.**
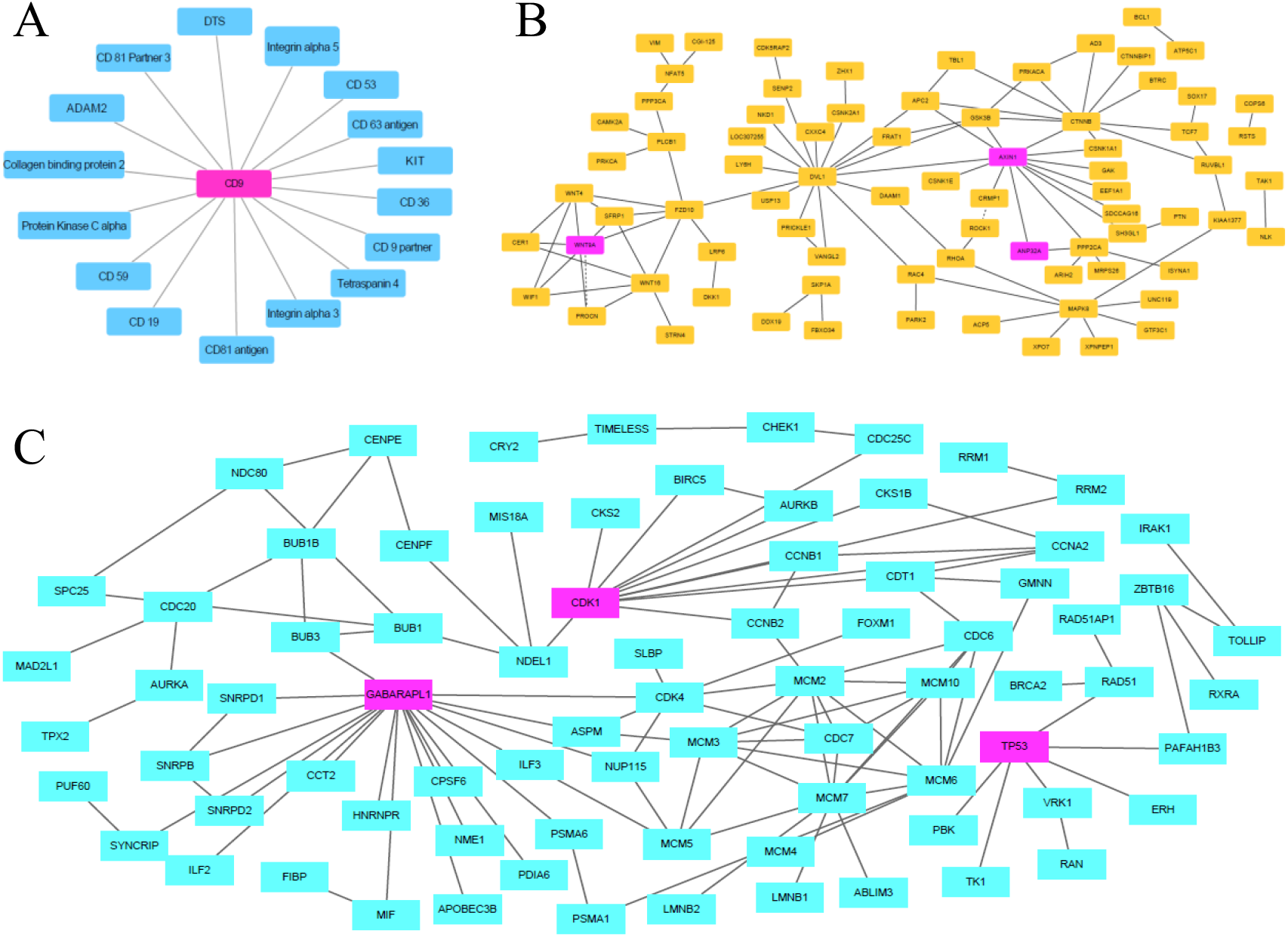
Predicted results on the three types PPIs networks. (A) Predicted results of PPIs networks of a one-core network for CD9. All 16 PPIs are truly predicted. (B) The predicted results of a crossover network, where WNT9A, CXXC4, AXIN1 and ANP32A are linked in the Wnt-related pathway. The solid lines are the interactions of true prediction, and the dotted lines are the interactions of false prediction. (C) Predicted results of PPIs networks of the cancer-specific differential genes. The network is composed of two components. The first component is marked in red and the second component is marked in blue. NDEL1 and GABARAPL1 connect the first component. TP53 is the main hub in the second component of this network. All 108 PPIs are truly predicted.

As shown in Fig. 6A, when using the GcForest-PPI model to predict a one-core network, all PPIs of the network are predicted successfully (16/16). The accuracy of GcForest-PPI is superior to Shen et al. (Shen, et al., 2007) and Ding et al. (Ding, Tang & Guo, 2016) (100% vs. 81.25%, 87.50%). CD9 plays a crucial role in sperm egg fusion, and myoblast fusion (Yang, et al., 2006). The palmitoylation of CD9 contributed to the interaction between CD81 and CD53 (Charrin, et al., 2002).

From Fig. 6B, we successfully predicted interacting protein pairs with accuracy of 97.92% on crossover network (94/96). GcForest-PPI is 21.88% higher than Shen et al. (Shen, et al., 2007) (97.92% vs. 76.04%). GcForest-PPI is 3.13% higher than that of Ding et al. (Ding, Tang & Guo, 2016) (97.92% vs. 94.79%). However, the relationship between ROCK1 and CRMP1 is not identified successfully. This maybe because ROCK1 is part of the noncanonical Wnt pathway, and GcForest-PPI is not very applicative to predict PPIs in this case. AXIN1 interacts with a variety of proteins and regulates multiple pathways (Luo & Lin, 2004). GcForest-PPI can truly predict the relationships between AXIN1 and its neighboring proteins. This means that the GcForest-PPI can be utilized to predict protein-protein signaling pathway networks, helping to gain insight into the significance of biology.

All PPIs in the cancer-specific network are successfully predicted (108/108). The cancer-specific network is composed of two sub-networks (Fig. 6C). The first sub-network is composed of 14 genes, where TP53 is the main hub. At the molecular level, TP53 is a gene associated with breast cancer (Andrysik, et al., 2017). The second subnetwork is a PPIs network consisting of 64 genes, where two down-regulated genes NDEL1 and GABARAPL1 link to two sub-modules (Wynne & Vallee, 2018). NDEL1 and LIS1 are essential for development, and they are thought to relate with cytoplasmic dynein (Hebbar, et al., 2008). NDEL1 contains phosphorylation sites for CDK1, CDK5 (Mori, et al., 2007). CDK5 phosphorylation of NDEL1 affects lysosome motility in axons, indicating CDK5 is important in cell growth and development (Klinman & Holzbaur, 2015; Pandey & Smith, 2011). NDEL1 is also closely related to the development of some diseases (Doobin, Kemal, Dantas & Vallee, 2016). All PPIs are predicted successfully on the cancer-specific network, indicating that the GcForest-PPI can provide new ideas to elucidate disease mechanisms, and design of new drugs.

## 4. Conclusion

With the rapid development of big data mining technology, the study of well-established computational predictive framework based on proteomics data is necessary. Using machine learning to automatically predict PPIs can provide reference for grasping disease pathogenesis, drug discovery and repositioning. We present a novel approach Gcforest-PPI for identifying PPIs, which uses PAAC, Auto, MMI, CTD, AAC-PSSM and DPC-PSSM to extract physicochemical features, sequence features and evolutionary features of PPIs. Then we use the elastic net to eliminate noise from extracted vectors, which could combine the advantages of L1 and L2 regularization and generate a sparse model and group effects. The comparison between raw features and optimal feature subset indicates the sequence information is more effective than physicochemical and evolutionary information when detecting PPIs. At the same time, deep forest is employed to predict PPIs for the first time, which uses XGBoost, RF and Extra-Trees to construct GcForest-PPI model. The cascade structure can mine nonlinear relationship to distinguish interacting and non-interacting samples. The results of *S. cerevisiae* and *H. pylori* indicate that GcForest-PPI can effectively identify PPIs. The prediction results of *C. elegans, E. coli, H. sapiens*, and *M. musculus* show that GcForest-PPI is capable of cross-species prediction and PPIs in *S. cerevisiae* include representation information of other species. Finally, the satisfactory scalability of the model is demonstrated by the one-core network, crossover network and cancer-specific network dataset, which can provide new ideas for exploring disease pathogenesis. In summary, GcForest-PPI can be a useful predictive tool for bioinformatics and proteomics.

Feature extraction from protein sequences is a key step based on machine learning. Although we combine the physicochemical and position information, sequence and composition information, and evolutionary information from primary interacting protein pairs, the comprehensive important features related to PPIs is still not elucidated. We are developing a python tool for feature extraction and feature selection to provide an online platform for the effective fusion of multiple feature information. DL has powerful representation learning ability and can mine more abstract essential features. Capsule neural network is a new deep learning framework. How to use capsule neural network is the next research direction.

## Acknowledgments

This work is supported by the National Nature Science Foundation of China (No. 61863010), the Key Research and Development Program of Shandong Province of China (2019GGX101001), the Natural Science Foundation of Shandong Province of China (Nos. ZR2017MA014 and ZR2018MC007), the Project of Shandong Province Higher Educational Science and Technology Program (No. J17KA159), and the Scientific Research Fund of Hunan Provincial Key Laboratory of Mathematical Modeling and Analysis in Engineering (No. 2018MMAEZD10). This work used the Extreme Science and Engineering Discovery Environment, which is supported by the National Science Foundation (No. ACI-1548562).

