## Supplementary Tables, Supplementary Figures for "Prediction of protein-protein interactions based on elastic net and deep forest"

**1. Supplementary Method Illustration**

File S1. Position specific score matrix.

File S2. Dimensionality reduction using the elastic net.

**2. Supplementary Tables**

Table S1. The values of the seven physicochemical properties for the 20 native amino acids.

Table S2. Amino acid physicochemical attributes and the division of the amino acids into three groups according to each attribute.

Table S3. Division of 20 amino acid types based on dipoles and volumes of side chains.

Table S4. Performance comparison with different $\lambda$ values on PPIs datasets.

Table S5. Performance comparison with different $lag$ values on PPIs datasets.

Table S6. Effect of selecting different penalty parameter $\alpha$ on the model performance.

Table S7. Effect of selecting different penalty parameter $\beta$ on the model performance.

Table S8. Comparison of prediction results on different dimensional reduction methods.

Table S9. The AUC and AUPR on different dimensional reduction methods.

Table S10. Prediction results of different classifiers on S. *cerevisiae*, *H. pylori* datasets.

Table S11. The AUC and AUPR of different classifiers on *S. cerevisiae*, *H. pylori* dataset.

**3. Supplementary Figures**

Fig. S1. 3-gram or 2-gram feature representation.

Fig. S2. The raw features and optimal features from different feature information.

**4. Supplementary References**

### 1. Supplementary Method Illustration

#### File S1. Position specific score matrix

In order to obtain the evolutionary information of the amino acid sequence, all the protein sequences in the dataset are compared with the non-redundant database SwissProt using the PSI-BLAST program (Altschul, et al., 1997). The program can search sequences based on the iterative BLAST search method. Evolutionary information in the position specific score matrix (PSSM) plays an important role in biological system analysis.

During the running process, the parameter E-value threshold of PSI-BLAST is set to 0.001, the maximum number of iterations is set to 3, and the remaining parameters are set by default. Then PSSM of each protein sequence is obtained. For a protein sequence whose length is $L$, PSSM is shown in equation (1).

$$P_{PSSM} = \begin{bmatrix} p_{1,1} & p_{1,2} & \cdots & p_{1,20} \\ \vdots & \vdots & \vdots & \vdots \\ p_{i,1} & p_{i,2} & \cdots & p_{i,20} \\ \vdots & \vdots & \vdots & \vdots \\ p_{L,1} & p_{L,2} & \cdots & p_{L,20} \end{bmatrix} \quad (1)$$

where each row of the PSSM represents a log likelihood score for amino acid substitutions occurring at corresponding positions in the query sequence. Where $P_{i,j}$ represents the $i$-th position of query sequence being mutated to type $j$ during evolution process. The scores are positive integers or negative integers. A positive integer indicates that more mutations have occurred in the alignment and a negative integer indicates that fewer substitutions have occurred in the alignment.

#### File S2. Dimensionality reduction using the elastic net

Although the fusion of physicochemical information, evolutional information, and sequence information of protein sequence can provide valuable initial feature information for predicting PPIs. The highly dimensional features will consume much calculation time and inevitably generate redundant and uncorrelated features, which may have a negative influence on PPIs prediction. The optimal feature subset of high-dimensional data can be obtained by the dimension reduction method, and the prediction model can be built up.

After the fusion of PseAAC, AD, MMI, CTD, AAC-PSSM, and DPC-PSSM, each protein sequence can generate a 1,074-dimensional feature vector. The protein pairs are transformed into a 2,148-dimension feature vector by concatenation. With the change of $\alpha$ and $\beta$, the performance of the model also changed. The larger the values of $\alpha$ and $\beta$ are, the more features are selected. The optimal parameters are determined via five-fold cross-validation. First, we fix $\beta$=0.1, taking $\alpha$ as 0.01, 0.02, 0.03, 0.04, and 0.05 and DF is employed as the classifier. It can be seen from Table S6, when $\alpha$=0.03, the ACC of the model achieves the maximum value. Secondly, we fix $\alpha$=0.03, and $\beta$ is taken as 0.1, 0.2, 0.3, 0.4, and 0.5. From

Table S7, when $\alpha$=0.03 and $\beta$=0.1, the *S. cerevisiae* and *H. pylori* datasets reach the peak value. Finally, the parameters of the elastic net are set $\alpha$=0.03 and $\beta$=0.1.

To further evaluate the advantages and disadvantages of the feature subset selected by the elastic net, we draw the original feature number and the optimal feature number of different feature information. As showcased in Fig. S2, PseAAC and AD characterize the physicochemical information of PPIs. MMI and CTD characterize the sequence information of PPIs. AAC-PSMM and DPC-PSSM characterize the evolutionary information of PPIs.

From Fig. S2, on the *S. cerevisiae* dataset, after dimension reduction of elastic net, physicochemical information features, sequence information features, and evolutionary information feature all exist in the selected optimal feature subsets. The sequence information accounts for 34.95% of its original features. And sequence information accounts for 57.56% of the total number of optimal features. In other words, the sequence information achieves the best performance for the prediction of PPIs, and physicochemical information is the next. On the *H. pylori* dataset, the optimal feature of sequence information accounts for 57.95% of the total number of optimal features, indicating that the MMI and CTD feature extraction methods can provide more critical feature information for PPI prediction. In summary, the six feature extraction methods of PseAAC, AD, MMI, CTD, AAC-PSSM, and DPC-PSSM obtain the physicochemical information, sequence information and evolutionary information. The elastic net can achieve the effective fusion of multiple information and eliminate redundancy. The optimal feature subset could characterize the feature information of PPIs.

### 2. Supplementary Tables

**Table S1**

The original values of the seven physicochemical properties for the 20 native amino acids.

| Amino acid | $\phi^{(1)}$ | $\phi^{(2)}$ | $\phi^{(3)}$ | $\phi^{(4)}$ | $\phi^{(5)}$ | $\phi^{(6)}$ | $\phi^{(7)}$ |
|---|---|---|---|---|---|---|---|
| A | 0.62 | -0.5 | 27.5 | 8.1 | 0.046 | 1.181 | 0.007187 |
| C | 0.29 | -1 | 44.6 | 5.5 | 0.128 | 1.461 | -0.03661 |
| D | -0.9 | 3 | 40 | 13 | 0.105 | 1.587 | -0.02382 |
| E | -0.74 | 3 | 62 | 12.3 | 0.151 | 1.862 | 0.006802 |
| F | 1.19 | -2.5 | 115.5 | 5.2 | 0.29 | 2.228 | 0.03755 |
| G | 0.48 | 0 | 0 | 9 | 0 | 0.881 | 0.1791 |
| H | -0.4 | -0.5 | 79 | 10.4 | 0.23 | 2.025 | -0.01069 |
| I | 1.38 | -1.8 | 93.5 | 5.2 | 0.186 | 1.81 | 0.02163 |
| K | -1.5 | 3 | 100 | 11.3 | 0.219 | 2.258 | 0.01771 |
| L | 1.06 | -1.8 | 93.5 | 4.9 | 0.186 | 1.931 | 0.05167 |
| M | 0.64 | -1.3 | 94.1 | 5.7 | 0.221 | 2.034 | 0.002683 |
| N | -0.78 | 2 | 58.7 | 11.6 | 0.134 | 1.655 | 0.005392 |
| P | 0.12 | 0 | 41.9 | 8 | 0.131 | 1.468 | 0.2395 |
| Q | -0.85 | 0.2 | 80.7 | 10.5 | 0.18 | 1.932 | 0.04921 |
| R | -2.53 | 3 | 105 | 10.5 | 0.291 | 2.56 | 0.04359 |
| S | -0.18 | 0.3 | 29.3 | 9.2 | 0.062 | 1.298 | 0.004627 |
| T | -0.05 | -0.4 | 51.3 | 8.6 | 0.108 | 1.525 | 0.003352 |
| V | 1.08 | -1.5 | 71.5 | 5.9 | 0.14 | 1.645 | 0.057 |
| W | 0.81 | -3.4 | 145.5 | 5.4 | 0.409 | 2.663 | 0.03798 |
| Y | 0.26 | -2.3 | 117.3 | 6.2 | 0.298 | 2.368 | 0.0236 |

*Note*: $\phi(1)$ is hydrophobicity; $\phi(2)$ is hydrophilicity; $\phi(3)$ is side chain volume; $\phi(4)$ is polarity; $\phi(5)$ is polarizability; $\phi(6)$ is solvent-accessible surface area and $\phi(7)$ is side chain net charge index.

**Table S2**

Amino acid physicochemical attributes and the division of the amino acids into three groups according to each attribute.

| Attribute | Division |
|---|---|
| Hydrophobicity_PRAM900101 | Polar: RKEDQN Neutral: GASTPHY Hydrophobicity: CLVIMFW |
| Hydrophobicity_ARGP820101 | Polar: QSTNGDE Neutral: RAHCKMV Hydrophobicity: LYPFIW |
| Hydrophobicity_ZIMJ680101 | Polar:QNGSWTDERA Neutral: HMCKV Hydrophobicity: LPFYI |
| Hydrophobicity_PONP930101 | Polar:KPDESNQT Neutral: GRHA Hydrophobicity: YMFWLCVI |
| Hydrophobicity_CASG920101 | Polar:KDEQPSRNTG Neutral: AHYMLV |

| | |
|---|---|
| | Hydrophobicity: FIWC |
| Hydrophobicity_ENGD860101 | Polar: RDKENQHYP Neutral :SGTAW Hydrophobicity: CVLIMF |
| Hydrophobicity_FASG890101 | Polar: KERSQD Neutral: NTPG Hydrophobicity: AYHWVMFLIC |
| Normalized van der Waals volume | Volume range:0-2.78 GASTPD<br>Volume range: 2.95-94.0 NVEQIL<br>Volume range: 4.03-8.08 MHKFRYW |
| Polarity | Polarity value: 4.9-6.2 LIFWCMVY<br>Polarity value: 8.0-9.2 PATGS<br>Polarity value: 10.4-13.0 HQRKNED |
| Polarizability | Polarizability value: 0-1.08 GASDT<br>Polarizability value: 0.128-120.186 GPNVEQIL<br>Polarizability value: 0.219-0.409 KMHFRYW |
| Charge | Positive: KR Neutral:ANCQGHILMFPSTWYV<br>Negative: D |
| Secondary structure | Helix:EALMQKRH Strand: VIYCWFT Coil: GNPSD |
| Solvent accessibility | Buried: ALFCGIVW Exposed: PKQEND<br>Intermediate: MPSTHY |

**Table S3**

Division of 20 amino acid types based on dipoles and volumes of side chains.

| Group 1 | Group 2 | Group 3 | Group 4 | Group 5 | Group 6 | Group 7 |
|---|---|---|---|---|---|---|
| A, G, V | C | D, E | F, I, L, P | H, N, Q, W | K, R | M, S, T, Y |

**Table S4**

Performance comparison with different $\lambda$ values on PPIs datasets.

| Dataset | Evaluation | $\lambda$ | | | | | |
|---|---|---|---|---|---|---|---|
| | | 1 | 3 | 5 | 7 | 9 | 11 |
| *S. cerevisiae* | ACC | 90.04 | 90.41 | 90.55 | 90.75 | 90.83 | **91.05** |
| | Recall | 88.36 | 88.54 | 88.58 | 88.72 | 88.92 | 89.08 |
| | Precision | 91.44 | 91.98 | 92.22 | 92.48 | 92.46 | 92.75 |
| | MCC | 0.8014 | 0.8088 | 0.8117 | 0.8157 | 0.8172 | 82.17 |
| *H. pylori* | ACC | 83.33 | 83.95 | 84.64 | 84.91 | 84.64 | **85.32** |
| | Recall | 85.11 | 85.46 | 85.53 | 85.80 | 84.91 | 85.60 |
| | Precision | 82.20 | 82.99 | 84.08 | 84.31 | 84.44 | 85.15 |
| | MCC | 66.74 | 67.97 | 69.33 | 69.85 | 69.29 | 70.67 |

**Table S5**

Performance comparison with different *lag* values on PPIs datasets.

| Dataset | Evaluation | *lag* | | | | | |
|---|---|---|---|---|---|---|---|
| | | 1 | 3 | 5 | 7 | 9 | 11 |
| | ACC | 91.60 | 93.60 | 93.81 | 93.93 | 93.94 | **94.10** |

| | | | | | | | |
|---|---|---|---|---|---|---|---|
| *S. cerevisiae* | Recall | 88.58 | 89.99 | 89.85 | 89.90 | 89.88 | 90.22 |
| | Precision | 94.28 | 97.00 | 97.58 | 97.79 | 97.83 | 97.82 |
| | MCC | 0.8336 | 0.8743 | 0.8789 | 0.8815 | 0.8818 | 0.8847 |
| *H. pylori* | ACC | 85.90 | 86.28 | **87.35** | 87.24 | 87.21 | 87.10 |
| | Recall | 81.82 | 82.58 | 84.16 | 84.50 | 83.61 | 84.77 |
| | Precision | 89.14 | 89.21 | 89.90 | 89.41 | 90.10 | 88.93 |
| | MCC | 0.7209 | 0.7281 | 0.7488 | 0.7464 | 0.7467 | 0.7432 |

**Table S6**

Effect of selecting different penalty parameter $\alpha$ on the model performance.

| Dataset | Evaluation | $\alpha$ | | | | |
|---|---|---|---|---|---|---|
| | | 0.01 | 0.02 | 0.03 | 0.04 | 0.05 |
| *S. cerevisiae* | ACC | 95.37 | 95.36 | **95.44** | 95.38 | 95.24 |
| | Recall | 92.72 | 92.60 | 92.72 | 92.97 | 92.40 |
| | Precision | 97.91 | 98.02 | 98.05 | 97.67 | 97.97 |
| | MCC | 0.9087 | 0.9086 | 0.9102 | 0.9086 | 0.9064 |
| *H. pylori* | ACC | 88.78 | 89.02 | **89.26** | 89.09 | **89.26** |
| | Recall | 89.85 | 89.44 | 89.71 | 89.50 | 90.19 |
| | Precision | 88.06 | 88.74 | 88.95 | 88.82 | 88.62 |
| | MCC | 0.7770 | 0.7809 | 0.7857 | 0.7825 | 0.7861 |

**Table S7**

Effect of selecting different penalty parameter $\beta$ on the model performance.

| Dataset | Evaluation | $\beta$ | | | | |
|---|---|---|---|---|---|---|
| | | 0.1 | 0.2 | 0.3 | 0.4 | 0.5 |
| *S. cerevisiae* | ACC | **95.44** | 95.17 | 95.18 | 95.11 | 94.87 |
| | Recall | 92.72 | 92.30 | 92.47 | 92.37 | 92.19 |
| | Precision | 98.05 | 97.94 | 97.77 | 97.74 | 97.42 |
| | MCC | 0.9102 | 0.9050 | 0.9050 | 0.9037 | 0.8987 |
| *H. pylori* | ACC | **89.26** | 89.20 | 88.48 | 87.72 | 87.76 |
| | Recall | 89.71 | 89.51 | 87.65 | 87.58 | 87.44 |
| | Precision | 88.95 | 89.00 | 89.14 | 87.90 | 88.08 |
| | MCC | 0.7857 | 0.7843 | 0.7700 | 0.7552 | 0.7563 |

**Table S8**

Comparison of prediction results on different dimensional reduction methods.

| Dataset | Evaluation | Method | | | | | | |
|---|---|---|---|---|---|---|---|---|
| | | EN | PCA | KPCA | LLE | SE | SVD | SSDR |
| *S. cerevisiae* | ACC | 95.44 | 89.81 | 88.57 | 86.98 | 84.81 | 90.87 | 77.54 |
| | Recall | 92.72 | 88.45 | 86.70 | 83.21 | 84.89 | 88.17 | 73.95 |
| | Precision | 98.05 | 90.97 | 90.08 | 90.01 | 84.78 | 93.23 | 79.71 |
| | MCC | 0.9102 | 0.7968 | 0.7720 | 0.7418 | 0.6964 | 81.88 | 55.25 |
| | ACC | 89.26 | 82.48 | 85.46 | 82.17 | 76.82 | 83.64 | 79.46 |

| | | | | | | | | |
|---|---|---|---|---|---|---|---|---|
| *H. pylori* | Recall | 89.71 | 78.46 | 82.44 | 81.28 | 77.43 | 79.35 | 73.87 |
| | Precision | 88.95 | 85.43 | 87.83 | 82.82 | 76.63 | 86.78 | 83.17 |
| | MCC | 0.7857 | 0.6530 | 0.7116 | 0.6442 | 0.5378 | 0.6758 | 0.5930 |

**Table S9**

The AUC and AUPR on different dimensional reduction methods.

| Dataset | Evaluation | Method | | | | | | |
|---|---|---|---|---|---|---|---|---|
| | | PCA | KPCA | LLE | SE | SVD | SSDR | EN |
| *S. cerevisiae* | AUC | 0.9603 | 0.9497 | 0.9302 | 0.9243 | 0.9664 | 0.8425 | 0.9864 |
| | AUPR | 0.9019 | 0.9230 | 0.8888 | 0.8509 | 0.9129 | 0.8560 | 0.9485 |
| *H. pylori* | AUC | 0.9545 | 0.9402 | 0.9088 | 0.9172 | 0.9605 | 0.7999 | 0.9816 |
| | AUPR | 0.8873 | 0.9209 | 0.8845 | 0.8356 | 0.9007 | 0.8254 | 0.9449 |

**Table S10**

Prediction results of different classifiers on S. *cerevisiae*, *H. pylori* dataset.

| Dataset | Model | ACC (%) | Recall (%) | Precision (%) | MCC |
|---|---|---|---|---|---|
| *S. cerevisiae* | LR | 85.79±0.72 | 84.47±0.94 | 86.77±0.84 | 0.7161±0.0144 |
| | NB | 71.75±1.17 | 63.67±1.40 | 75.99±1.98 | 0.4410±0.0248 |
| | KNN | 89.19±0.58 | 83.98±0.23 | 93.75±1.06 | 0.788±0.0124 |
| | AdaBoost | 92.37±0.77 | 90.15±0.84 | 94.33±0.80 | 0.8482±0.0153 |
| | RF | 93.17±0.46 | 89.24±0.74 | 96.86±0.73 | 0.8662±0.0092 |
| | SVM | 90.61±0.61 | 89.61±0.71 | 91.43±0.79 | 0.8123±0.0123 |
| | GcForest | 95.44±0.18 | 92.72±0.44 | 98.05±0.25 | 0.9102±0.0035 |
| *H. pylori* | LR | 82.75±1.48 | 84.09±1.86 | 81.90±1.41 | 0.6553±0.0297 |
| | NB | 68.28±2.41 | 87.45±3.26 | 63.22±1.79 | 0.3964±0.0520 |
| | KNN | 72.43±1.19 | 95.88±1.96 | 65.26±0.72 | 0.5083±0.0305 |
| | AdaBoost | 85.22±1.12 | 86.97±0.96 | 84.06±1.64 | 0.705±0.0219 |
| | RF | 86.35±1.24 | 87.52±2.15 | 85.52±0.90 | 0.7274±0.0249 |
| | SVM | 84.71±1.36 | 83.54±1.91 | 85.56±1.62 | 0.6945±0.0273 |
| | GcForest | 89.26±1.07 | 89.71±2.26 | 88.95±1.36 | 0.7857±0.0212 |

**Table S11**

The AUC and AUPR of different classifiers on S. *cerevisiae*, *H. pylori* dataset

| Dataset | Evaluation | Method | | | | | | |
|---|---|---|---|---|---|---|---|---|
| | | LR | NB | KNN | AdaBoost | RF | SVM | GcForest |
| *S. cerevisiae* | AUC | 0.9298 | 0.7914 | 0.9503 | 0.9750 | 0.9762 | 0.9653 | 0.9864 |
| | AUPR | 0.8996 | 0.8022 | 0.8722 | 0.9225 | 0.9427 | 0.9264 | 0.9485 |
| *H. pylori* | AUC | 0.9189 | 0.7721 | 0.9270 | 0.9693 | 0.9704 | 0.9616 | 0.9816 |
| | AUPR | 0.9091 | 0.7653 | 0.8764 | 0.9229 | 0.9447 | 0.9246 | 0.9449 |

### 3. Supplementary Figures

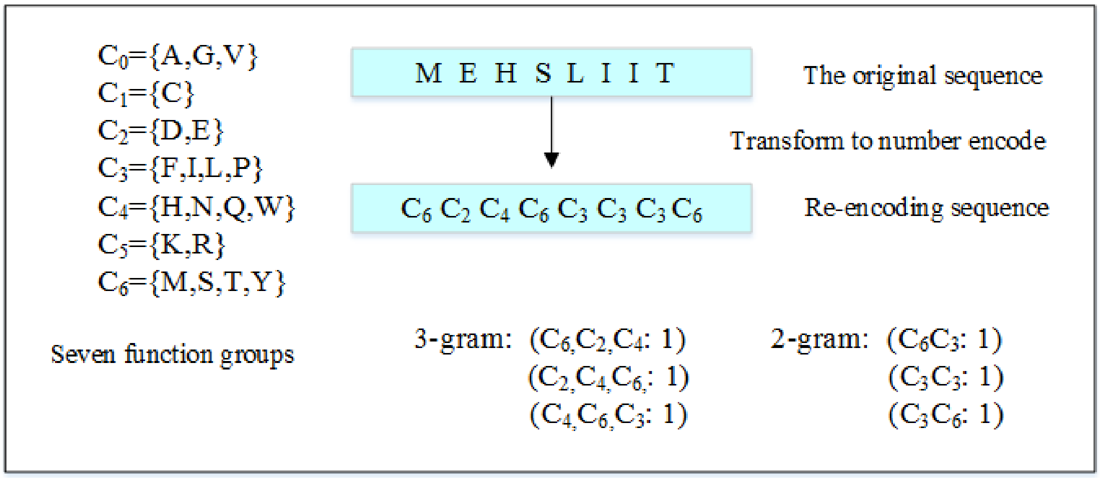


**Fig. S1.** 3-gram or 2-gram feature representation.

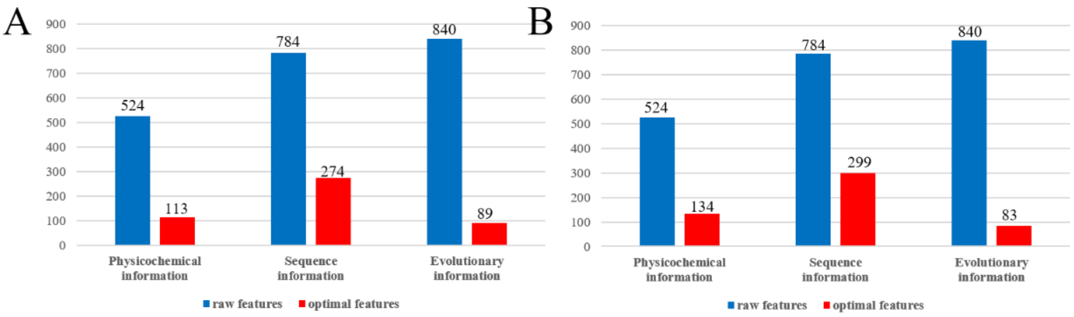


**Fig. S2.** The raw features and optimal features from different feature information. (A) The raw features and optimal features of *S. cerevisiae* dataset. (B) The raw features and optimal features on *H. pylori* data set.
